# Next generation AMA1-based plasmids for enhanced heterologous expression in filamentous fungi

**DOI:** 10.1101/2024.06.01.596972

**Authors:** Indra Roux, Clara Woodcraft, Nicolau Sbaraini, Amy Pepper, Emily Wong, Joe Bracegirdle, Yit-Heng Chooi

## Abstract

Episomal AMA1-based plasmids are increasingly used for expressing biosynthetic pathways and CRISPR/Cas systems in filamentous fungi cell factories due to their high transformation efficiency and multicopy nature. However, gene expression from AMA1 plasmids has been observed to be highly heterogeneous in growing mycelia. To overcome this limitation, here we developed next-generation AMA1-based plasmids that ensure homogenous and strong expression. We achieved this by evaluating various degradation tags fused to the auxotrophic marker gene on the AMA1 plasmid, which introduces a more stringent selection pressure throughout multicellular fungal growth. With these improved plasmids, we observed in *Aspergillus nidulans* a 5-fold increase in the expression of a fluorescent reporter, a doubling in the efficiency of a CRISPRa system for genome mining, and a up to a 10-fold increase in the production of heterologous natural product metabolites. This strategy has the potential to be applied to diverse filamentous fungi.

## Introduction

Filamentous fungi play key roles in biotechnology, serving as versatile cell factories for producing industrial enzymes, organic acids, and bioactive small molecules. Fungal mycelium also has applications in biomaterials, bioremediation, biomass processing and microbial foods (Maini Rekdal et al., 2024; Meyer et al., 2020). Notably, *Aspergillus* stands as the most popular genus in the academic literature and patent landscape for fungal biotechnology (Füting et al., 2021).

Moreover, the fungal kingdom remains an untapped reservoir of pharmaceutical lead compounds, such as polyketides, non-ribosomal peptides and other natural product small molecules. These are encoded in biosynthetic gene clusters (BGCs) often spanning >20 kb (Robey et al., 2021). Fungal genomes also encode a diverse array of unexplored enzymes with potential biorefinery applications such as carbohydrate-active enzymes (CAZymes). This broad biosynthetic and enzymatic diversity is observed both at inter- and intraspecies levels (Robey et al., 2021; Vesth et al., 2018). Promising cryptic genes for enzyme discovery are often found in fungal taxa which are challenging to cultivate or genetically intractable. One of the strategies to functionally characterise candidate enzymes is expressing them in a heterologous host, with *A. nidulans* emerging as a popular chassis for genome mining (Caesar et al., 2020).

The expression of heterologous genes or BGCs in a filamentous fungi host is usually achieved through chromosomal integration or maintenance in self-replicating extrachromosomal plasmids, such as those containing the AMA1 replicator. Chromosomal expression offers genetic stability, but random gene integration often requires extensive screening to identify productive strains due to locus-specific and pleiotropic effects (Lubertozzi and Keasling, 2009). Moreover, as the efficiency of targeted homologous recombination decreases with insert size, multiple transformation rounds are required to integrate large BGCs (Chiang et al., 2013). Although genetic tools such CRISPR or Cre-lox recombinase can enhance efficiency, single copy integration often results in low production yields when compared to AMA1-based expression (Roux and Chooi, 2022a; Woodcraft et al., 2022).

In contrast, AMA1-based episomal plasmids offer advantages such as high transformation efficiency and increased yields due to the multi-copy nature of heterologous genes (Aleksenko and Clutterbuck, 1997). They are particularly suitable for expressing long BGCs in *Aspergillus* and *Penicillum* hosts for natural product discovery (Caesar et al., 2023; Roux et al., 2021; Sbaraini et al., 2021). For example, AMA1-plasmids have been used to construct fungal genomic libraries with insert sizes of ∼100 kb to screen BGCs in an *Aspergillus* host (Bok et al., 2015; Clevenger et al., 2017). They also facilitate the elucidation of biosynthesis by enabling combinatorial expression of gene sets from plasmids with different selection markers (Chiang et al., 2022; Roux and Chooi, 2022b). Furthermore, AMA1-plasmids have been widely used across diverse fungi to express synthetic biology tools, such as CRISPR/Cas components for gene editing (Katayama et al., 2019; Nødvig et al., 2018; Woodcraft et al., 2022) and to modulate gene expression via CRISPRa (Mózsik et al., 2021a; Roux et al., 2020) or RNAi constructs (Shimizu et al., 2015). Notably, AMA1-based plasmids have been recently used for in vivo assembly in *Aspergillus*, bypassing the need for prior cloning in *E. coli* (Jarczynska et al., 2022).

While the AMA1 replicator sequence was originally isolated from an *A. nidulans* genomic library (Aleksenko and Clutterbuck, 1997), it has become an integral part of the genetic toolbox for diverse filamentous fungi from the Eurotiomycetes order, including *Aspergillus* (e.g., *A. oryzae, A. niger, A. aculeatus)* (Jarczynska et al., 2021; Katayama et al., 2019), *Penicillium* (Mózsik et al., 2021b) and *Talaromyces* (Nielsen et al., 2017). Interestingly, AMA1 plasmids have also been shown to sustain replication in distantly related mushroom-forming Sordariomycetes (Kanematsu and Shimizu, 2015; Meng et al., 2022) and aid gene delivery in Basidiomycetes (Xie et al., 2022).

However, a major drawback of expressing genes from episomal plasmids is the requirement for continuous selection pressure to ensure plasmid maintenance. Some AMA1-based plasmids such as pPTRII by Takara Bio, rely on antifungal resistance selection markers (e.g., *ptrA, hygR*)(Kubodera et al., 2002; Meng et al., 2022). However, for stable bioproduction, a cost-effective and scalable strategy based on auxotrophic markers (e.g., *pyrG, argB, riboB, pyroA*) were implemented in *E. coli*-yeast-fungi shuttle vectors, such as pKW20088, pYFAC or pYT, which are widely used for natural product genome mining (Chiang et al., 2022; Roux and Chooi, 2022b; Tsunematsu et al., 2013; Yan et al., 2018). Auxotrophic marker genes, most commonly the pyrimidine pathway gene *pyrG*, is used in engineered fungal strains for plasmid selection by complementation when grown on selective media. Yet, AMA1-based expression can be heterogeneous and unstable over time, even with different markers (Aleksenko and Clutterbuck, 1997; Roux and Chooi, 2022a; Shimizu et al., 2015). This instability could be potentially attributed to the multicellular nature of filamentous growth and the capability to transport resources along hyphae (Pantazopoulou and Diallinas, 2007). Thus, we reasoned that the genetic and phenotypic stability of AMA1-based plasmids might be compromised by the occurrence of “cheating” hyphal cells, which escape selection pressure by inheriting selection maker proteins from other plasmid-carrying cells in the mycelium. Furthermore, excess prototrophy-conferring metabolites could be secreted by the plasmid-carrying cells into the media. This potential lack of stringent selection pressure could also lead to a decrease in plasmid copy number.

To overcome this limitation, we drew inspiration from studies conducted in baker’s yeast where plasmid copy number was increased by limiting the expression of the marker gene (Chen et al., 2012; Lian et al., 2016). We aimed to investigate the effect of constraining the expression of the auxotrophic marker gene in the AMA1 plasmid during multicellular fungal growth as means to increase productivity. Here, we develop a set of next generation AMA1-based plasmids based on the popular auxotrophic marker *pyrG*. We increased plasmid selection stringency by fusing the selection marker to a degradation tag to decrease protein stability and limit its potential spread. We demonstrate that AMA1-based plasmids containing a degron-tagged marker outperformed those harbouring the parental untagged marker for the expression of fluorescent reporter proteins, delivery of guide RNA for CRISPR activation and heterologous natural product metabolite production.

## Results

### AMA1-based plasmids with stringent selection markers support homogenous and high fluorescent protein expression

To enhance the phenotypic stability of extrachromosomal AMA1-based expression in filamentous fungi, we aimed to develop next-generation plasmids that impose more rigorous selection pressure. We reasoned that by increasing the turnover of selection marker proteins, we could reduce their effective expression and intercellular carry over. This would minimise the numbers of potential cheating cells loosing AMA1 plasmid in the growing mycelia. To achieve this, we used as a starting point the AMA1-based shuttle plasmid pYFAC, which also supports replication in *E. coli* and *S. cerevisiae* for cloning (Chooi et al., 2017). This plasmid contains the selection marker *pyrG*, which encodes the orotidine-5′-decarboxylase from *A. fumigatus*, and is used paired with auxotroph strains, in our case *A. nidulans* LO8030 (Chiang et al., 2016), grown on selective minimal media (no uracil or uridine supplementation).

One of the popular degrons in eukaryotic synthetic biology applications are the bipartite ubiquitin N-terminal (ubi-N) degrons (Houser et al., 2012). In this system, an N-terminal ubiquitin moiety is cleaved to reveal a new amino acid directing proteins for degradation via N-end rule pathway (Bachmair and Varshavsky, 1989; Passmore et al., 1986). To our knowledge, the synthetic ubiquitin-N degrons (ubi-N) had not yet been employed in filamentous fungi as a genetic tool. We constructed two new plasmid versions by fusing the N-terminus of *pyrG* to ubi-N degrons derived from the versions optimised by Houser et al (2012) including the flexible linker Δk (Supporting Text 1). In these constructs the 76 amino acid ubiquitin is derived from yeast, whose sequence is conserved to ubiquitin moieties native from *A. nidulans*. We built two ubi-N degrons: one with a highly destabilising tyrosine residue (*ubi-Y-pyrG*) and another with less destabilising methionine (*ubi-M-pyrG*) (Houser et al., 2012). Additionally, protein motifs directly targeted by the proteosome can be used as degradation tag, as the PEST motif from ornithine decarboxylase (ODC). This motif has been demonstrated to act as a degradation tag fused to fluorescent proteins in Penicillum spp. (Pohl, 2020). Thus, we additionally evaluated this degron to constrain the expression of the *pyrG* marker (*ODC-pyrG*).

To evaluate the impact of the different selection markers in the phenotypic stability of AMA1-based expression, we constitutively expressed the fluorescent reporter mCherry from the various plasmid versions (Figure 1A). When analysing the transformant colonies on selective solid agar medium by fluorescent photography, we observed a strong increase in fluorescence in the strains expressing mCherry from plasmids with *ubi-Y-pyrG* and *ubi-M-pyrG* markers, compared to expression from the original untagged *pyrG* plasmid (Figure 1B, Figure S2). These colonies also showed a more homogeneous fluorescence compared with the patchy fluorescence on *pyrG* colonies on solid media. To quantitatively investigate mCherry expression, we examined spores from different transformant colonies by flow cytometry (Figure 1C, Figure S1A). As we had previously observed (Roux and Chooi, 2022a) the AMA1-based plasmid with untagged *pyrG* presents a wide bimodal distribution of mCherry signal, with most of the spores being in the lower end of the fluorescence signal. In contrast, the spores harbouring plasmids with the markers *ubi-Y-pyrG* and *ubi-M-pyrG* present a unimodal distribution located in the higher end of classic *pyrG* plasmid signal, representing up to a 5-fold increase in mean fluorescence (Figure 1C). When comparing between these two new plasmids, the mean fluorescence of *ubi-Y-pyrG* colonies (98±9 AU) was much higher and robust than *ubi-M-pyrG* colonies (mean 53±23 AU), while both significantly higher than the mean of colonies with untagged *pyrG* (18±6 AU). On the other hand, the colonies expressing mCherry from the *ODC-pyrG* plasmid showed patchy fluorescence and only marginal increases in mean fluorescence (mean 28±5 AU) (Figure S1B).

**Figure 1.**
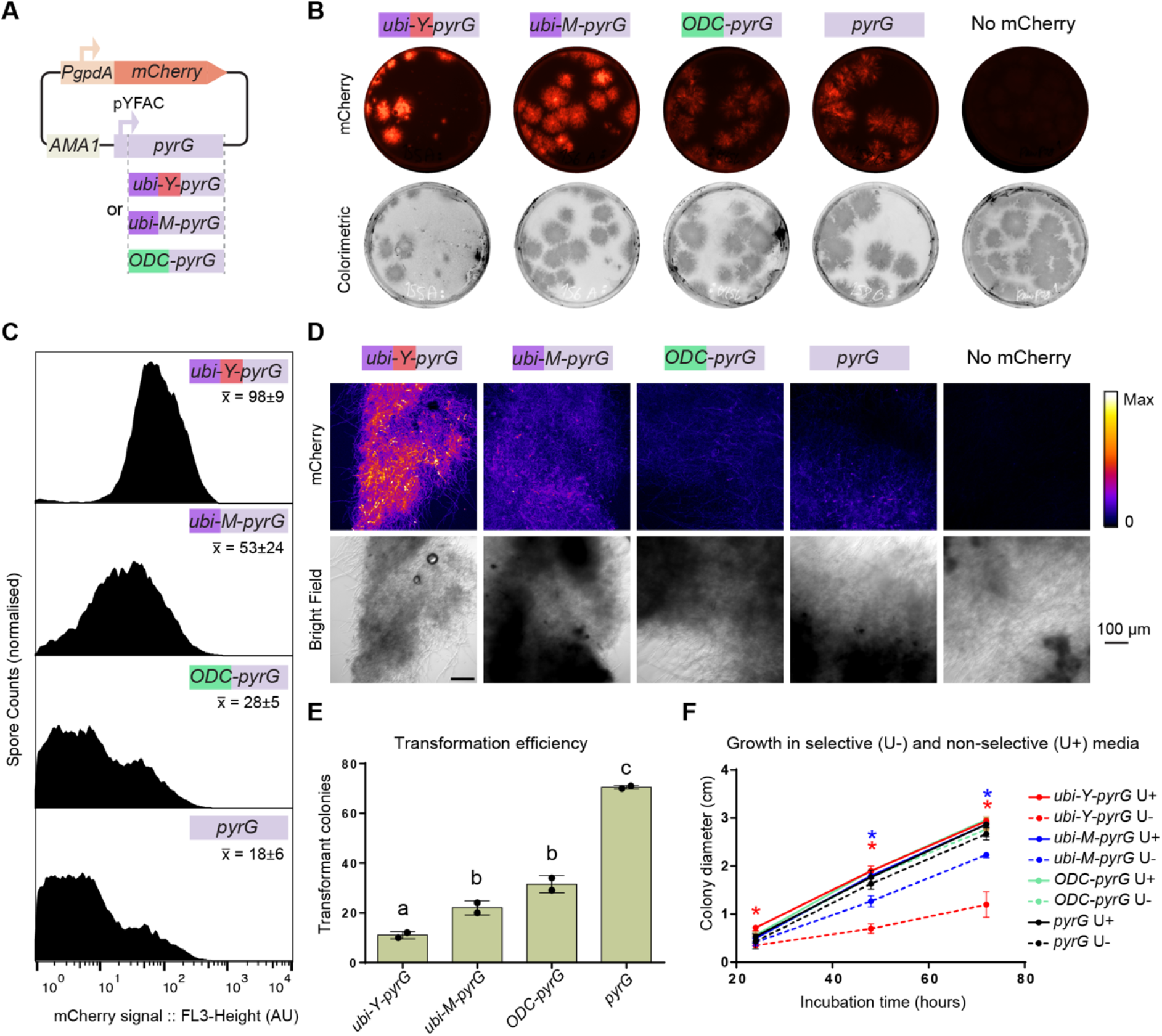
Enhanced expression of mCherry fluorescent protein using next-generation AMA1-based plasmids. **A**. Schematic of the AMA1 plasmids analysed for mCherry expression, with different variants of the selection marker. **B** Fluorescent photographs of transformant *Aspergillus nidulans* colonies reveal increased fluorescence in colonies transformed with *ubi-M-pyrG* and *ubi-Y-pyrG* plasmids. **C** Flow cytometry analysis of mCherry fluorescence in spores from a transformant colony show that the most uniform and high mCherry signal is achieved with the *ubi-Y-pyrG* plasmid. Mean fluorescent between replicates from different transformant colonies indicated, and histogram of replicates in Figure S1A. **D** Confocal microscopy images of mycelia grown in liquid culture show increased expression in mycelia containing *ubi-M-pyrG* and *ubi-Y-pyrG* plasmids. ImageJ FIRE calibration bar represents different levels of mCherry signal. Replicates in Fig. S3. **E**. Evaluation of the transformation efficiency for different plasmids at equimolar plasmid concentrations shows decreased transformation efficiency for plasmids with *pyrG* fused to degradation tags. Letters denote significantly different groups as determined by ANOVA with post hoc Tukey’s test. **F**. Comparison of colony growth rates on solid media under selected and non-selected conditions indicates slower growth for strains carrying *ubi-M-pyrG* and *ubi-Y-pyrG* plasmids under selective conditions. Asterisks represent Padj<0.05 by Welch’s T-test for differences in diameter between selective and non-selective media for strains carrying each plasmid.

As fungal bioprocesses often take place in submerged liquid culture, we also analysed mycelia grown overnight in selective liquid media by confocal microscopy. In this setting, *ubi-Y-pyrG* strains showed a higher increase in fluorescence compared to mycelia from *pyrG* strains (Figure 1D, Figure S3). Overall, *ubi-Y-pyrG* AMA1 plasmid stands the best candidate for heterologous expression based on the intensity of the fluorescent reporter expression in solid and liquid culture.

Besides the improvement in expression level, we also noticed differences in transformation efficiency and growth rate among colonies carrying different plasmids. When transforming at equal molar concentrations, we observed a decrease in the transformation efficiency of all three new plasmids compared to the untagged *pyrG* plasmid (Figure 1E). Furthermore, colonies transformed with the *ubi-Y-pyrG* and *ubi-M-pyrG* containing plasmids showed a significant decrease in their radial growth rates on solid media under selective conditions (Figure 1F). This suggests that in solid media lacking supplements, the limited effective expression of Ubi-Y-pyrG and Ubi-M-pyrG impose a constraint in growth. This is not observed with the untagged *pyrG*, where the growth rate in selective and non-selective conditions is indistinguishable (Figure 1F). This inverse relation between robust mCherry expression and colony growth rate may indicate that the stringent selective markers limit the growth of “escape” or “cheater” mycelia that complements Δ*pyrG* without requiring maintenance of the plasmid in growing mycelia.

### CRISPRa efficiency is optimised by delivery of gRNA from stringent AMA1-based plasmids

Synthetic biology tools such as CRISPR devices are commonly delivered on AMA1-based plasmids in filamentous fungi (Woodcraft et al., 2022). Some of these tools rely on stable expression through fungal growth, such as CRISPR-based activators (CRISPRa). In CRISPRa, a nuclease-deactivated Cas protein fused to a transcriptional activation domain acts as a synthetic transcription factor that can be programmed to activate the expression of silent or lowly expressed genes. This has multiple potential applications in fungi, such as activating silent biosynthetic gene clusters for secondary metabolite discovery. As the programmable CRISPR guide RNAs (gRNAs) can be short lived (Zhang et al., 2023), this system is particularly vulnerable to cells losing AMA1-based gRNA expression. Therefore, we aimed to improve the efficiency of CRISPRa by delivering gRNAs from the AMA1-based plasmids with degron-tagged markers. To test this, we built next-generation plasmids for gRNA delivery as an expansion to our previously developed CRISPR/dCas12a-VPR CRISPRa toolkit for genome mining of cryptic natural product genes (Roux et al., 2020). To ensure independence to Cas protein expression, we used an *A. nidulans* strain with constitutive chromosomally expressed dCas12a-VPR. As a reporter, we tested the activation of the endogenous chromosomal *A. nidulans* nonribosomal peptide synthetase-like (NRPS-like) *micA* by measuring the metabolite product microperfuranone (Figure 2A). The strains harbouring gRNA delivery from AMA1 plasmids with degron-tagged markers *ubi-Y-pyrG* and *ubi-M-pyrG* yielded a significant two-fold increase in microperfuranone titre than the untagged *pyrG* plasmid control (Figure 2B). In contrast, no significant increase in production was observed from the gRNA expression plasmid with *ODC-pyrG*, and thus we do not pursue further experiments with *ODC-pyrG* plasmids.

**Figure 2.**
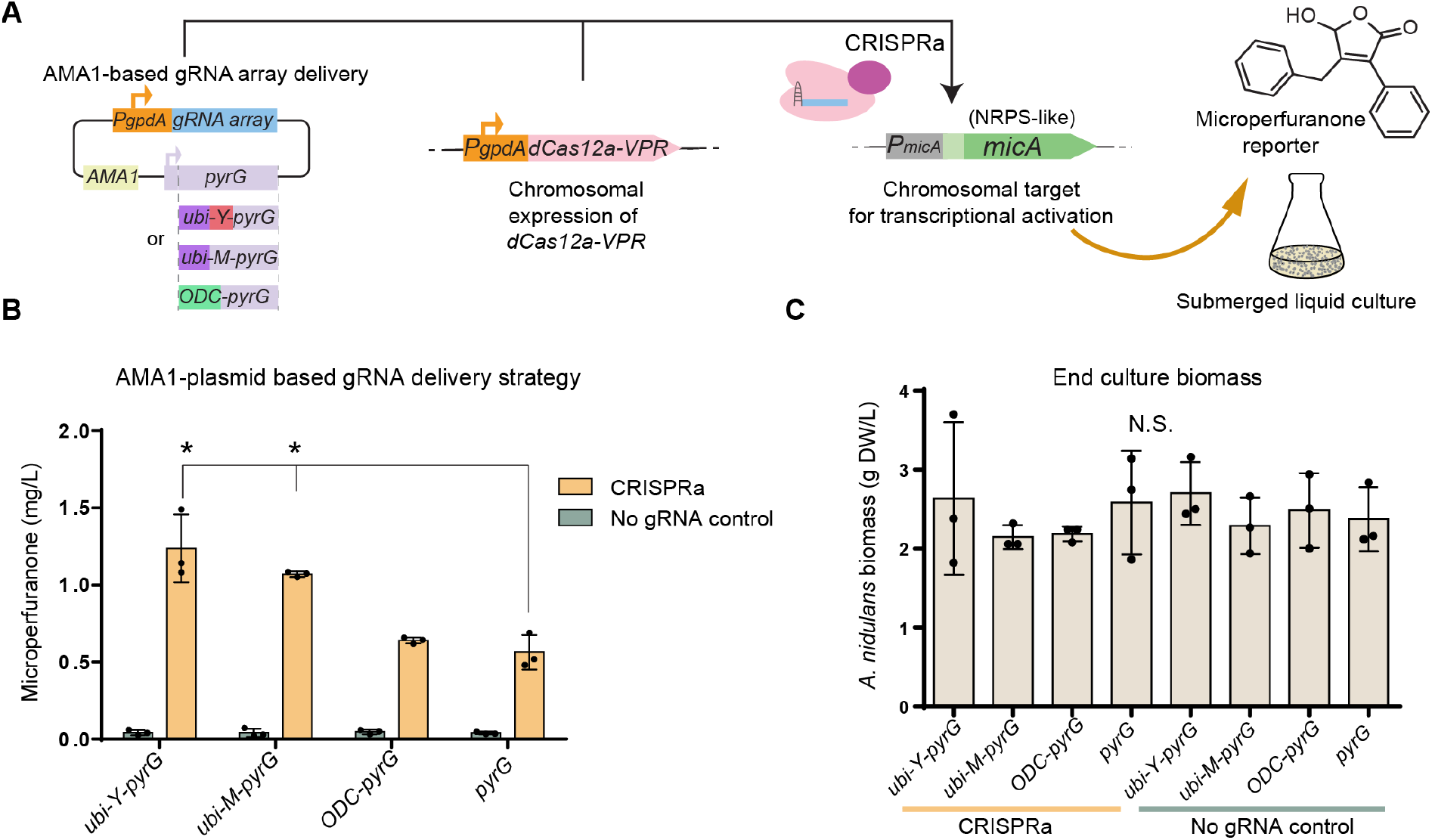
Improved delivery of gRNA for CRISPRa using *ubi-Y/M-pyrG* plasmids results in enhanced transcriptional activation, inferred by increased metabolite production. **A**. Experimental design, evaluating the delivery of CRISPR gRNAs from different variants of AMA1 episomal plasmids by measuring increases in the production of microperfuranone, encoded by the target biosynthetic gene *micA*. **B**. The CRISPRa-mediated output microperfuranone is increased by a 2-fold in the strains expressing gRNA arrays from *ubi-Y/M-pyrG* plasmids. Asterisks indicate significant difference by Welch’s t-test (P-adj <0.05). **C**. No significant changes were observed in end culture biomass between *Aspergillus nidulans* strains harbouring the different AMA1 expression plasmids. Dots represent 3 biological replicates.

To investigate the possible impact of the AMA1 plasmids with different degron-tagged and untagged *pyrG* marker on mycelial growth during liquid culture, we quantified the dry weight of the biomass at the end of culturing. Surprisingly, we observed no significant difference in the end point biomass between cultures harbouring *ubi-Y-pyrG, ubi-M-pyrG, ODC-pyrG* and untagged *pyrG* plasmids for either functional CRISPRa or no gRNA control strains (Figure 2C), although we cannot exclude the possibility of differences in growth dynamics.

### Improving episomal production of heterologous secondary metabolites

Increasing the yields and robustness of heterologous secondary metabolite production is of major interest for higher throughput natural product discovery. To evaluate the capability of AMA1 plasmids with degron-tagged markers to enhance secondary metabolite production, we cloned the NRPS-like gene *micA* with its native promoter P_*micA*_ into *ubi-Y-pyrG* and *ubi-M-pyrG* plasmids (Figure 3A). To mimic conditions for scalable high throughput screening, we cultured the strains in 1.5 mL of media in deep-well plates. Notably, both *ubi-Y/M* degron-tagged *pyrG* plasmids expressing P_*micA*_*-micA* showed a significant peak matching microperfuranone in the chromatogram, while the untagged *pyrG* plasmid showed only a minor peak close to the noise level (Figure 3B). Upon quantification, the strains with the *ubi-Y-pyrG* plasmid version supported a remarkable 20-fold increase in microperfuranone production to the untagged *pyrG* plasmid, while a 5-fold increase was observed for *ubi-M-pyrG* (Figure 3B). This highlights the utility of the AMA1 plasmids with degron-tagged *pyrG* marker to facilitate small volume culture at increased yields, which can enhance throughput for secondary metabolite detection by culturing in deep-well plate format.

**Figure 3.**
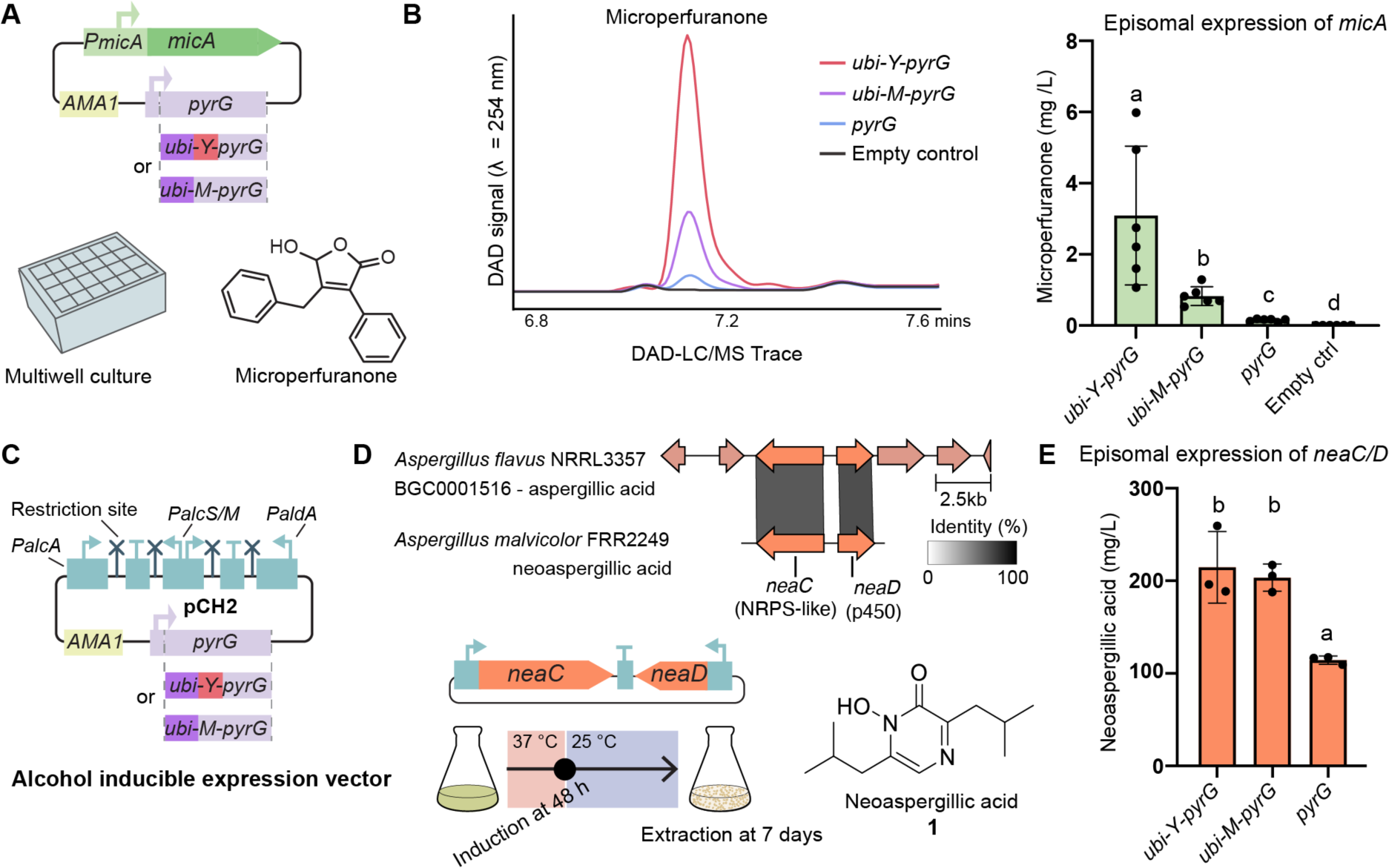
Increased yield of heterologous fungal secondary metabolites with degron-tagged AMA1 plasmids. **A**. Schematic of the plasmids evaluated for microperfuranone expression, in strains cultivated in multiwell plates. **B**. Representative DAD-LC/MS chromatogram showing the differences in the peak of microperfuranone, and respective quantification shown as bar plot. The dots represent six biological replicates, letters represent significantly different groups by Welch’s t-test (P-adj <0.05). **C**. Schematic of the four-promoter alcohol inducible expression plasmid pCH2, now expanded to the marker genes *ubi-Y-pyrG* and *ubi-M-pyrG*. **D**. Aspergillic acid biosynthetic gene cluster from *Aspergillus flavus*, and homologue NRPS-like (*neaC*) and p450 (*neaD*) genes from *Aspergillus malvicolor* characterised in this work. Grayscale bars linking genes indicate the extent of amino acid identity of the encoded protein (0% transparent, 100% black). Schematic of the expression plasmid and inducible culturing strategy for neoaspergillic acid (**1**) below. **E**. Yields of neoaspergillic acid. The dots represent 3 biological replicates. One-way ANOVA with posthoc Tukey’s test was performed.

To make the plasmids with *ubi-Y-pyrG* and *ubi-M-pyrG* markers more useful for heterologous expression of cryptic BGCs, we integrated the degron-tagged markers with the previously reported pYFAC-CH2 plasmid (Figure 3C). In this system, the expression of up to four genes is driven by a series of alcohol inducible promoters (P_*alcA*_, P_*alcS/M*_, *and* P_*aldA*_), which enables separating the biomass growth from compound production (Hu et al., 2019; Roux and Chooi, 2022b). To validate the newly generated plasmids (pYFAC-CH2-ubi-Y-pyrG and pYFAC-CH2-ubi-M-pyrG), we aimed to express an uncharacterised BGC from the *Aspergillus malvicolor* strain FRR2249, which displays regions of homology to a previously characterised aspergillic acid gene cluster in *A. flavus* (MIBIG BGC0001516) (Figure 3D, Figure S4). For the biosynthesis of aspergillic acid or the isomer neoaspergillic acid, two genes are necessary: the NRPS-like enzyme *neaC (asaC)* and the cytochrome P450 gene *neaD (asaD)* (Lebar et al., 2022, 2018). Therefore, *A. malvicolor*-derived *neaC* and *neaD* were cloned into alcohol inducible promoters in the new degron-tagged *pyrG* plasmids, as well as into the original plasmid (pYFAC-CH2).

After culturing and induction of transformant strains (Fig. 3D), we observe a new peak in all strains expressing *neaC* and *neaD* (Fig. S4). Subsequent large-scale cultures followed by extraction and purification, yielded approximately 50 mg of compound **1** (Fig S4) as a yellow solid. The molecular formula of compound **1** was inferred as C_12_H_20_N_2_O_2_ based on the protonated ion peak at m/z 225.1593 (calcd 225.1598, Δ – 0.42 ppm) in the (+) HR-ESIMS spectrum. The 1D ^1^H NMR spectrum gave evidence for one olefin, two methines, two methylenes and four methyl groups, while the ^13^C spectrum coupled with the multiplicity-edited HSQC gave additional evidence for these as well as an additional three sp2 hybridised carbons, one of which was an amide (Figure S5–7, Table S4). From literature searches, it was clear compound **1** was either aspergillic acid or an isomer (Dunn et al., 1949; Yuan et al., 2023). When correlations in COSY and HMBC spectra were analysed to define the side chain spin systems, it was observed that both contained chemically equivalent methyl groups that correlated to a methine and then onto a methylene, as is the R-group of leucine. Therefore compound **1** was identified as neoaspergillic acid (Figure 3C), which has been recently characterised as a candidate antifungal against *Candida albicans*, as well as exhibiting antibacterial and antitumoral activities (Yuan et al., 2023).

To evaluate the impact of different AMA1 plasmids on neoaspergillic acid yields, we compared productivity in 50 mL shake flask cultures. Transformants harbouring *neaCD* on the *ubi-Y-pyrG* and *ubi-M-pyrG* plasmids almost doubled in yield for neoaspergillic acid compared to transformants with *neaCD* on untagged *pyrG* plasmid (Figure 3D). The yields of neoaspergillic acid from transformants with degron-tagged *pyrG* plasmids reached around 200 mg/L without any optimisation (Figure 3C), highlighting that the new degron-tagged *pyrG* plasmid system can increase the yield even in notably productive expression constructs.

## Discussion

AMA1 plasmids have been game-changing in filamentous fungi genetics, enabling the popularisation of CRISPR tools, accelerating biosynthesis studies, and propelling synthetic biology development (Caesar et al., 2020; Woodcraft et al., 2022). Click or tap here to enter text.However, gene expression on AMA1 plasmids can be unstable and heterogeneous, which compromises their performance and scalability. To address this challenge and improve the utility of AMA1-based plasmids, here we developed a set of next-generation AMA1 plasmids featuring stringent auxotrophic selection marker genes that are fused to degron tags (*ubi-Y-pyrG* and *ubi-M-pyrG*) to ensure robust and homogenous phenotypic stability in fungal culture. Across all the scenarios tested in this study, the *ubi-Y-pyrG* and *ubi-M-pyrG* plasmids outperformed the AMA1 plasmids with untagged *pyrG*. We demonstrated that this new set of plasmids can improve natural product yields for scaled up compound isolation, as well as enabling efficient miniaturised culturing for higher throughput natural product screening. Importantly, even the most stringent degradation tag, the system maintains reliable transformation efficiency with >10 transformant colonies per miniaturised transformation in microtubes, which supports genome mining of enzymes and metabolic pathways.

Our initial test on fluorescent protein expression revealed *ubi-Y-pyrG* as an outstanding candidate for heterologous expression, although we observed a cost in fungal growth rate on solid media. Nevertheless, in liquid media we did not observe a significant impact on biomass production with either *ubi-Y-pyrG* or *ubi-M-pyrG* plasmids. In the case for CRISPR gRNA delivery, both systems performed comparable with a 2-fold increase in efficiency to the untagged *pyrG* plasmid. This is likely because once gRNA content is saturated, no increases in CRISPRa efficiency are expected. In the case of secondary metabolite production, both *ubi-Y-pyrG* and *ubi-M-pyrG* plasmids performed comparably, with the yield of neoaspergillic acid double the level obtained with the untagged *pyrG* plasmid. In the case of microperfuranone production in small volume deep-well cultures, *ubi-Y-pyrG* almost quadrupled the yields of *ubi-M-pyrG* strains, reaching a 10-fold increase compared to *pyrG* plasmid. Overall, *ubi-Y-pyrG* stands as the best performer plasmid for heterologous expression, with *ubi-M-pyrG* as an alternative with less impact in transformation efficiency and growth rate on solid medium.

Additionally, we characterised *A. malvicolor neaC* and *neaD* as genes responsible for neoaspergillic acid biosynthesis. The degron-tagged AMA1-plasmids achieved a yield of 200 mg/L, which is higher than the yields of neoaspergillic acid in the literature with engineered strains (140 mg/L in *A. melleus* Δ*mcrA*) (Yuan et al., 2023). This highlights the power of next generation AMA1-based heterologous expression in *A. nidulans* for bioprospecting and secondary metabolite production.

This toolkit has great potential to be portable to other fungal taxa where AMA1 functions as a replicative plasmid. While this work centred on the popular *pyrG* selection marker as it is widely used across fungal taxa, the strategy of N-degron tagging could be likely applied to other auxotrophic markers or antifungal resistance genes Alternatively, constraining marker gene expression using weak promoters could be evaluated as this was effective to increase copy numbers in plasmids with the antifungal resistance gene hygromycin in yeast (Lian et al., 2016).

While the aim of this work is developing a system for increased productivity, the possible mechanisms behind auxotrophic plasmid stability in filamentous fungi remain to be explored. Auxotrophic laboratory strains of baker yeast harbouring episomal plasmids selected with auxotrophic markers tend to yield a subpopulation where plasmid loss is compensated by cross-feeding (Yu et al., 2022). Further studies documenting the extent of cross feeding in filamentous fungi could facilitate the rational design of auxotrophic markers in future works.

In conclusion, the next-generation AMA1 plasmids here developed can help accelerate filamentous fungi genetics and synthetic biology for enzyme and natural product discovery. The increased phenotypic stability and robust performance between transformant colonies can help maximise throughput and versatility in fungal biotechnology.

## Experimental procedures

### Plasmid construction

A list of plasmids used is found in Table S1 and all oligonucleotides used in Table S2, relevant plasmids are being deposited in Addgene. Plasmids pYFAC-ubi-Y-pyrG (Addgene ID# 184497), pYFAC-ubi-M-pyrG (Addgene ID# 184498) and pYFAC-ODC-pyrG were built by isothermal assembly using NEBuilder HiFi DNA Assembly Master Mix (New England Biolabs, MA, USA). First, pYFAC (Addgene ID# 168982)(Chooi et al., 2017) was digested with NotI and MscI to release the original *pyrG* marker. Then, in-frame fusions of *pyrG* and degradation tags were created by isothermal assembly. The sequence of *ubi-Y* and *ubi-M* were respectively amplified from the plasmids pB2GW7-ubi-Y-GFP and pB2GW7-ubi-M-GFP, which were a gift from Brendan Kidd (UWA), and contain degradation tags derived from plasmids from Houser et al (2012), Click or tap here to enter text.which are derived from *S. cerevisiae* with its original codon usage. The sequence of the degron from ornithine-decarboxylase (ODC) was based on sequences tested in *Penicillium chrysogenum* by Pohl et al (Pohl, 2020), and cloned embedded in oligonucleotides. *PgpdA-mCherry* expression cassette was amplified from pRecomb-loxP-mCherry-4Gcloningsite-lox2272-66 (Addgene ID# 168787). The gRNA expression cassette was amplified from pCRI040 which contains four Cas12a crRNAs targeting upstream of *micA* (Roux et al., 2020). *micA* was amplified from *A. nidulans* genomic DNA. The plasmid for heterologous expression with alcohol inducible promoters pYFAC-CH2-ubi-Y-pyrG (Addgene ID# 221131) and pYFAC-CH2-ubi-M-pyrG (Addgene ID# 221132) incorporate the four-promoter expression cassette from pYFAC-CH2 (Addgene ID #168978) (Hu et al., 2019). These two plasmids were built using yeast-mediated homologous recombination as previously described (Gietz and Schiestl, 2007; Roux and Chooi, 2022b) combining PCR-amplified fragments and digested fragments (Lebar et al., 2018). The genes *neaC* (Protein ID XAV45317.1) and *neaD* (Protein ID XAV45316.1) for neoaspergillic acid production were identified from the genome of *A. malvicolor* strain FRR2249, which we deposited in GenBank under accession number PP792998. The *A. malvicolor* strain was a gift from Microbial Screening Technologies (Australia). These genes were PCR-amplified from gDNA and cloned into the PacI and NotI sites of pYFAC-CH2-ubi-Y-pyrG and pYFAC-CH2-ubi-M-pyrG using yeast-mediated homologous recombination.

### *Aspergillus nidulans* transformation and culturing

A list of all strains built, and parental strains are found in Table S3, all derived from *A. nidulans* LO8030 which has several BGCs removed for minimal metabolite background (Δ*nku, pyroA4, riboB2, pyrG89*) (Chiang et al., 2016). Fungal protoplast transformation was performed using the PEG-CaCl_2_ method as previously described (Roux and Chooi, 2022b) and grown for 4 days at 37 °C in Glucose Minimal Media (GMM) supplemented with riboflavin and pyridoxine (See Supporting Methods). When evaluating the transformation efficiency, 4.5 μg of plasmid quantified by Qubit dsDNA broad range (Invitrogen) was used to transform 10^7^ *A. nidulans* protoplasts. This DNA amount was derived from what is usually obtained with 5 μl of plasmid miniprep, which is in the range of maximum DNA volume recommended for miniaturised protoplast transformations. Similar amounts were used in all the other transformation events described.

To evaluate the growth rate in solid media, 2 μl of spore solution (10^5^ spores/mL) was inoculated in the centre of selective and non-selective GMM plates. Colony diameter was measured during 3 days of growth at 37 °C. To evaluate effects in growth on liquid media, biomass dry weight at the end of liquid culture was obtained by filtering mycelia with preweighted filter paper and drying in a 50 °C oven before final weighting.

### Fluorescence microscopy, flow cytometry and fluorescent photography

For fluorescence photography, plates grown four days since transformation at 37 °C were analyzed in ChemiDoc MP Imaging System (Bio-Rad) and images were recorded with ImageLab software V6.1.0 (Bio-Rad). MCherry images were obtained using the excitation source Green Epi illumination and the emission filter 605/50 nm with an exposure time of 0.5 seconds. Bright field was captured with white illumination and a standard emission filter in automatic exposure.

In the flow cytometry analysis, spores were collected in water with 0.01% Tween and filtered through a syringe containing sterile cotton to remove residual mycelia and diluted in water to a concentration of ∼10^6^ spores/mL. Data acquisition was performed using a FACSCalibur (BD Biosciences) flow cytometer operated with filtered water as sheath fluid. mCherry signal was observed with a 488 nm excitation laser and the filter FL3 (≥670 nm) and 35,000 events were recorded per sample or events within 4 minutes of run time as collected for the filtered water control. The data was processed using FlowJo V10 software (TreeStar). Examples of gating are included in Figure S1C.

For fluorescent confocal microscopy, mycelia were grown overnight at 37 °C in selective liquid GMM and washed with 0.6 M MgSO_4_ before mounting in the slides. Images were taken with a Nikon C2 Confocal Microscope using 488 excitation and 640 nm filter for mCherry, and all figures were imaged using the same settings collecting a stack of 10 in ∼50 μm. The software Fiji ImageJ was used for Z projection using sum slices. Mcherry channel is represented with ImageJ FIRE scale for ease of interpretation.

### Secondary metabolite analysis by LC-DAD-MS

For microperfuranone production analysis as activated by CRISPRa, analysis was performed as previously described (Roux et al., 2020). Strains were cultured in 50 mL of selective GMM in 250 mL flasks for 72 hours at 37 °C with shaking at 200 rpm. For episomal expression of *micA*, the reactions were scaled down to a deep well microtiter plate of 24 × 11mL wells (17 × 17 × 40 mm) with 1.5 mL GMM per well and cultivated at 37 °C with 200 rpm shaking. Following three days of incubation, the culture was extracted by 1:1 liquid-liquid partitioning with ethyl acetate, methanol, acetic acid mix (89.5: 10 : 0.5). The crude extracts were dried down under N_2_ flow and redissolved in 150 μL of methanol for LC-DAD-MS analysis. Figure source data is found in Supporting File 2.

For neoaspergillic acid small scale cultures, *A. nidulans* spores were inoculated (∼10^8^/L) into 50 mL of liquid selective GMM media in 250 mL flasks. The cultures were grown at 37 °C with shaking at 180 rpm for 120 hours. Following this, 3 mL/L of methyl ethyl ketone was added to induce the expression of *Alc* promoters. Cultures were then incubated at 25 °C with shaking at 180 rpm for 48 hours. The supernatant was isolated by vacuum filtration and 10 mL were extracted employing an organic solvent mixture as described above. The crude extracts were dried down under a N_2_ flow and re-dissolved in 1 mL methanol for LC-DAD-MS analysis.

Metabolite profile analyses of the *A. nidulans* cultures harbouring the constructed plasmids, as well as the control plasmids, were performed using an Agilent 1260 liquid chromatography (LC) system, coupled to a diode array detector (DAD) and an Agilent 6130 Quadrupole mass spectrometer (MS) with an ESI source. Chromatographic separation was carried out at 40 °C, employing a Kinetex C18 column (2.6 μm, 2.1 × 100 mm; Phenomenex). Chromatographic separation was achieved with a linear gradient of 5–95% acetonitrile-H2O (0.1% (v/v) formic acid) over 7 minutes, followed by 95% acetonitrile for 3 minutes, with a flow rate of 0.6 mL/min. The MS data were collected in the m/z range of 100–1000. A standard curve was prepared by injecting 5, 4, 3, 2, and 1 μg of purified neoaspergillic acid, for quantification, the integrated Peak Area were recorded, employing three biological replicates with three technical replicates each (Supporting File 2). Microperfuranone was quantified as previously described (Roux et al., 2020).

### Neoaspergillic acid isolation and NMR structure elucidation

For isolation of neoaspergillic acid, *A. nidulans* strain LO8030 harbouring harbouring neaCD was grown in 1 L shake-flask culture of liquid GMM medium supplemented with riboflavin and pyridoxine following the induction regime previously described. Supernatant extraction was performed as described in the above section for microperfuranone. After initial LC-DAD-MS evaluation, the crude extract was purified using an Agilent 1260 Infinity II LC system, coupled to a DAD. Chromatographic separation was carried out on a preparative C18 (5 μm, 150 × 21.2 mm; Agilent) column, with separation achieved over a linear gradient of 10– 70% acetonitrile-H2O (0.1% (v/v) trifluoroacetic acid) over 20 minutes, at a flow rate of 12 mL/min. NMR spectra were recorded in 5 mm Pyrex tubes (Wilmad, USA) on a Bruker Avance III HD 600 spectrometer. All NMR spectra were obtained at 25 °C and processed using Bruker Topspin 3.5 software.

### Statistics

Statistical analyses were conducted with GraphPad Prism (v 8.3.0) or R (v 4.2.3). Differences between compound production were determined by one-way ANOVA with posthoc Tukey’s test (*p* < 0.05) or by Two-sided Welch’s T-test with BH multiplicity correction, as indicated in figure captions. The letters above bars indicate statistically different groups.

## Acknowledgements

We thank Dr Brendan Kidd (UWA) for helpful discussion and the plasmids pB2GW7-ubi-Y-GFP and pB2GW7-ubi-M-GFP. We thank Genomics WA for whole plasmid sequencing and Guy Ben-Ari from UWA Cell Biology and Imaging Research Infrastructure Platform for advice on confocal microscopy. We thank Microbial Screening Technology for providing the *A. malvicolor* strain FRR2249. We thank Dr. Cameron L.M. Gilchrist for his assistance with genome assembly. *A. nidulans* LO8030 is a gift from Berl Oakley. This work was supported by the Australian Research Council (DP210102180 and FT160100233). C.W. is the recipient of a UWA PhD Scholarship. We thank UWA’s Separation Science & Mass Spectrometry Platform, and the UWA’s Centre for Microscopy Characterisation and Analysis (CMCA) Additionally, we are grateful for the assistance provided by the Future Health Research and Innovation Infrastructure Fund (FHRIFGCOVID19/14).

## Author contribution

I.R. conceptualised the project and lead in writing the manuscript. I.R., C.W. and N.S. designed and conducted experiments and analysis. A.P. and E.W. performed experiments. J.B. performed NMR analysis. Y.H.C supervised the project and contributed to the final manuscript. All authors discussed the results and commented on the manuscript.

