## Supporting Figures, Tables and Text for "Next generation AMA1-based plasmids for enhanced heterologous expression in filamentous fungi"

### Supporting methods

#### Culture media

Glucose minimal medium (GMM). For 1 L, 10 g Glucose, 6g $NaNO_3$, 1.52 g $K_2HPO_4$, 0.52 g KCl, 0.52 g $MgSO_4 \cdot 7H_2O$, 22 mg $ZnSO_4 \cdot 7H_2O$, 11 mg $H_3BO_3$, 5 mg $MnCl_2 \cdot 4H2O$, 1.6 mg $FeSO_4 \cdot 7H_2O$, 1.6 mg $CoCl_2 \cdot 5H_2O$, 1.6 mg $CuSO_4 \cdot 5H_2O$, 1.1 mg $(NH_4)_6Mo_7O_{24} \cdot 4H_2O$, 50 mg Na4EDTA, media adjusted to pH 6.5). Supplements added when required, 10 mL/L of riboflavin 0.025% w/v, 1 mL/L of pyridoxine-HCl 0.2% w/v, or uridine 1.26 g/L and uracil 0.56 g/L.

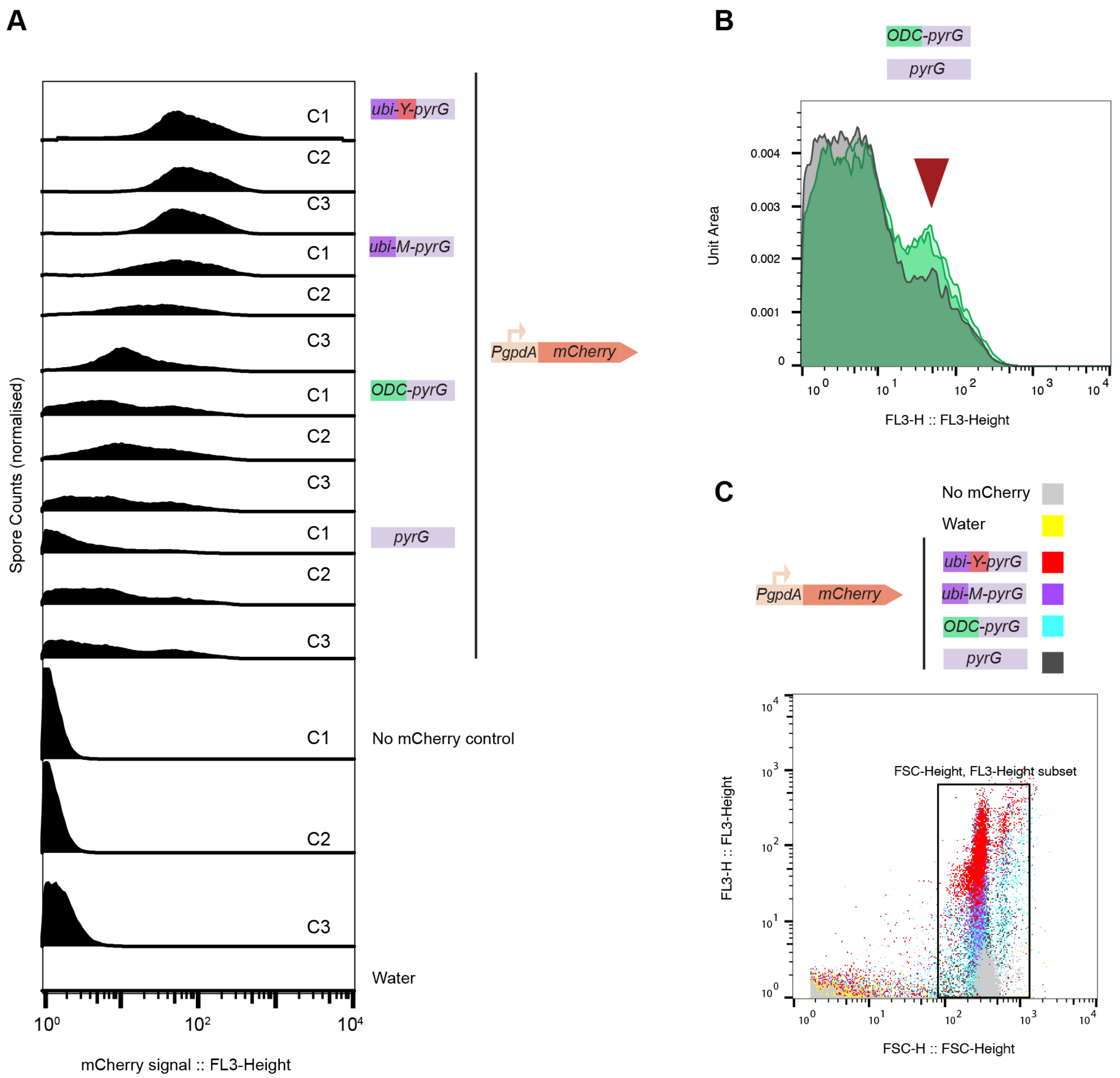

**Figure S1. Flow cytometry of spores from colonies expressing mCherry from AMA1 plasmids with different *pyrG* marker versions. A.** Biological replicates of colonies from Figure 1, showing a consistent increase in the fluorescent intensity of the tags *ubi-Y-pyrG* and *ubi-M-pyrG*. **B.** Overlay of the ODC-*pyrG* tag and *pyrG* mCherry normalised fluorescence histogram. A minimal increase of ≈6% population is observed in most colonies of ODC-*pyrG* (red arrow). **C.** Gating strategy used in the flow cytometry analysis.

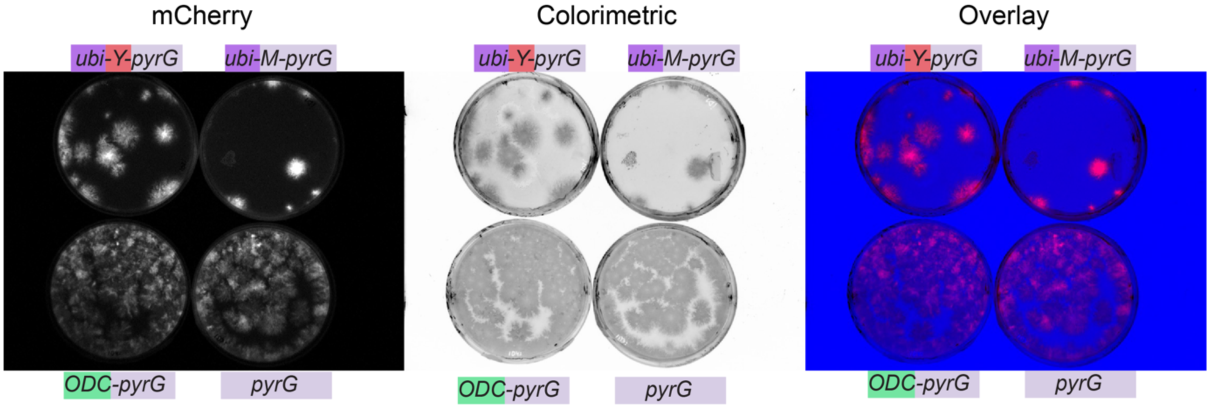

**Figure S2. Fluorescence photography of mCherry expressing mCherry from AMA1 plasmids with different *pyrG* marker versions.** This is an independent biological replicate from Figure 1. No differences are observed between ODC-pyrG and pyrG transformant plates.

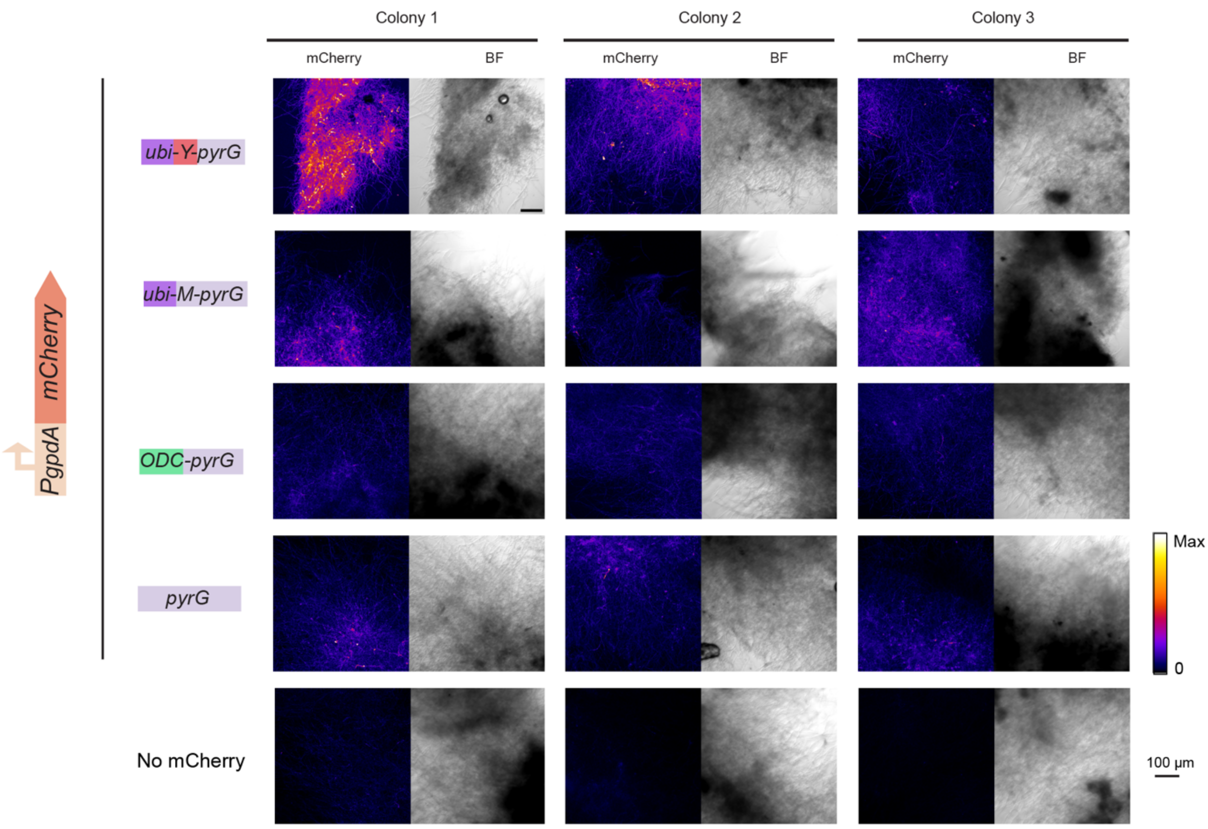

**Figure S3. Confocal microscopy of mycelia grown in liquid culture shows different levels of mCherry expression.** Three biological replicates were analysed from separate transformant colonies. Calibration bar represents different levels of mCherry signal as per ImageJ FIRE scale.

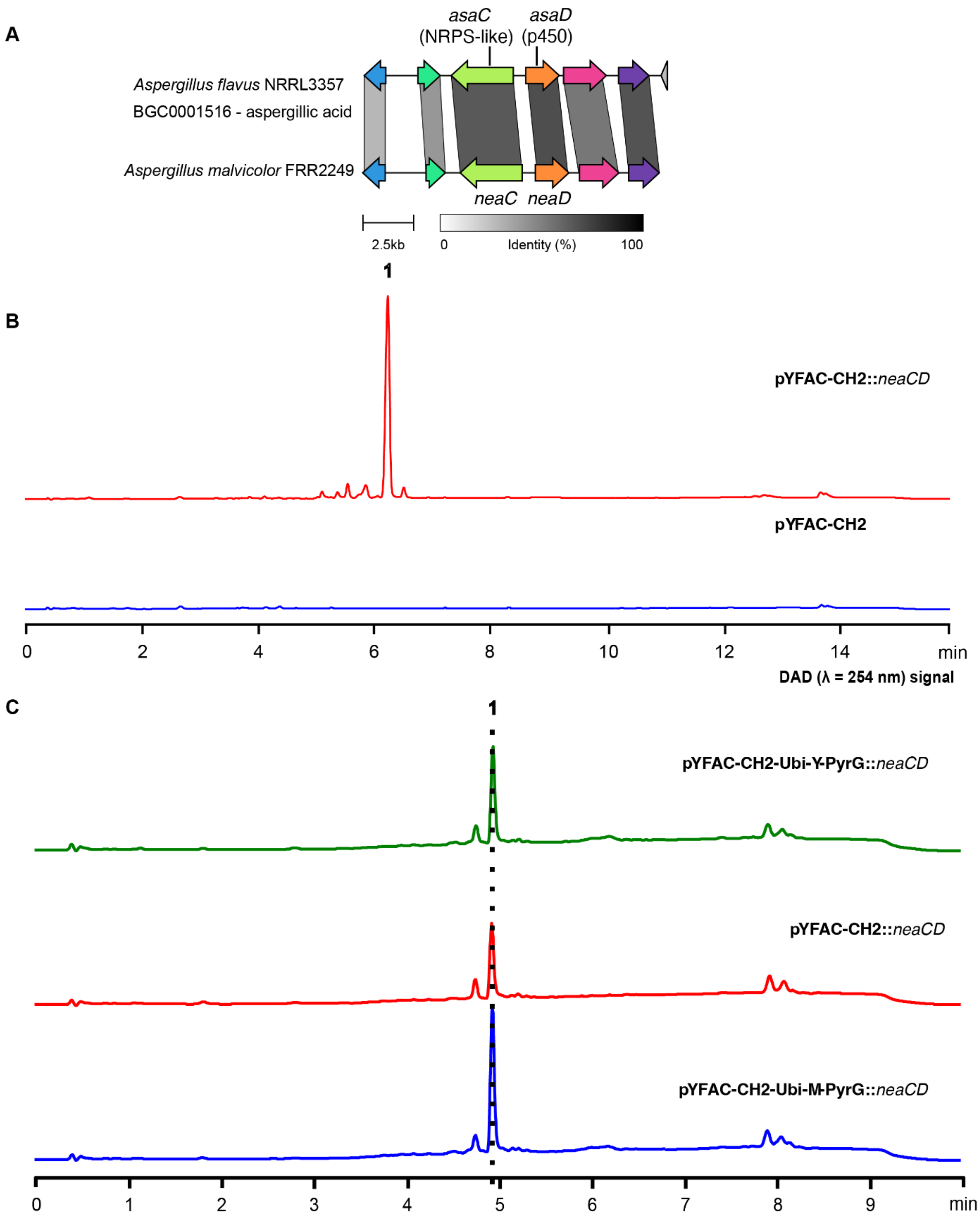

**Figure S4. *Aspergillus malvicolor neaCD* characterisation and LC-DAD profiles. A.** Homology between *Aspergillus flavus* aspergillic acid biosynthetic gene cluster and the *Aspergillus malvicolor* neoaspergillic acid biosynthetic gene cluster. Protein identity is represented in grey scale as by clinker. **B.** LC-DAD chromatogram at 254 nm shows **1** (identified as neoaspergillic acid) as the main product of *neaCD*. **C.** Chromatograms showing the increase in the production of **1** with *ubi-Y-pyrG* and *ubi-M-pyrG* plasmids, compared to the untagged *pyrG* plasmid (pFAC-CH2).

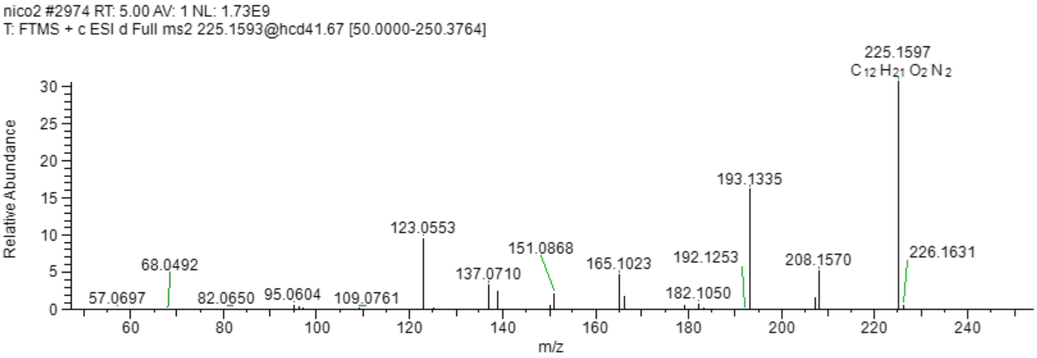

**Figure S5. HR-ESIMS spectrum of 1.**

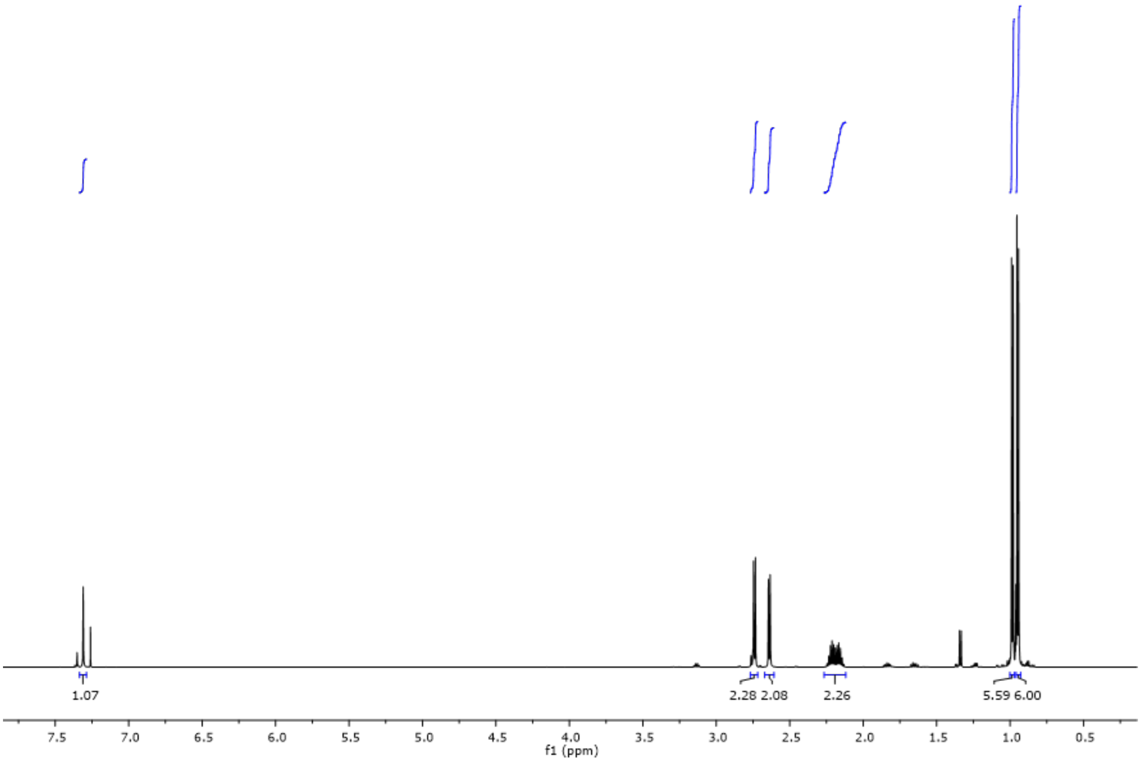

**Figure S6. $^1$H NMR (600 MHz) spectrum of 1 in $CDCl_3$.**

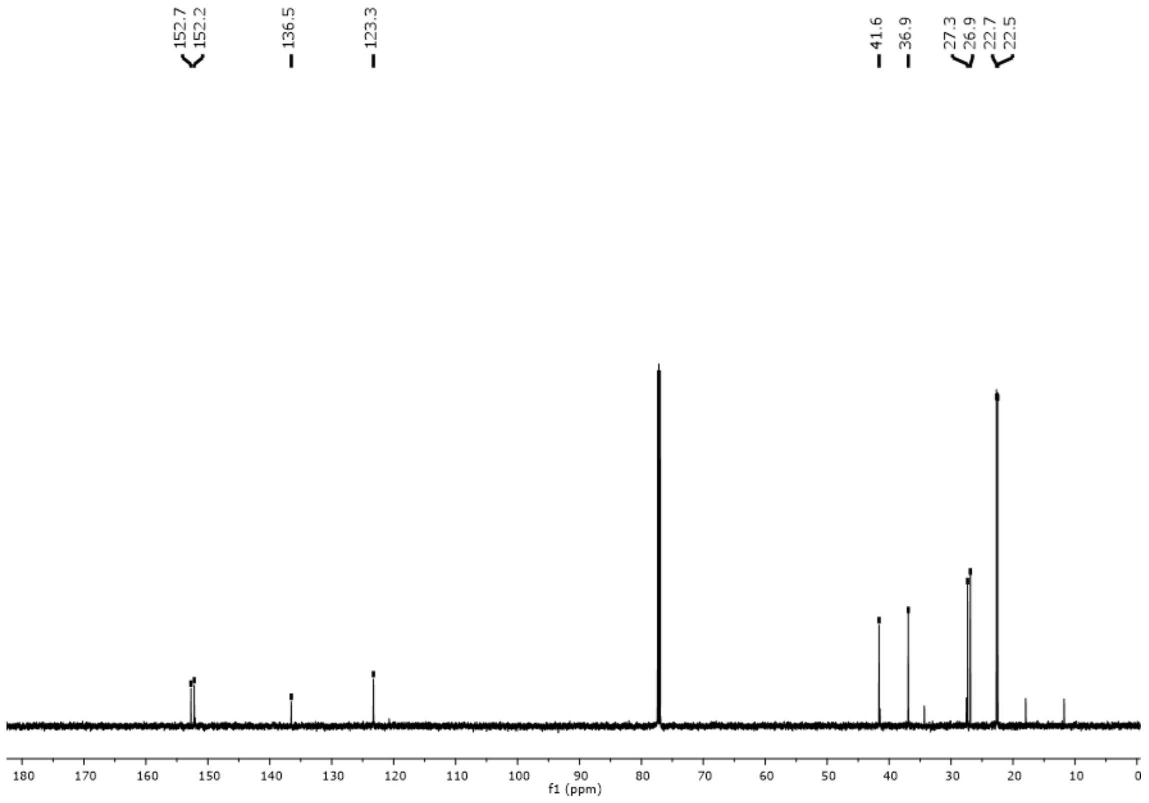

**Figure S7. $^{13}C$ NMR (150 MHz) spectrum of 1 in $CDCl_3$.**

**Table S1 Plasmids used in this study**

| Vector name | Vector purpose | Origin | Size (kb) | Addgene ID |
|---|---|---|---|---|
| pYFAC-ubi-Y-pyrG | AMA1-based backbone vector with *ubi-Y-pyrG* marker. | This work | 14.5 | 184497 |
| pYFAC-ubi-M-pyrG | AMA1-based backbone vector with *ubi-M-pyrG* marker. | This work | 14.5 | 184498 |
| pYFAC-ODC -pyrG | AMA1-based backbone vector with *ODC-pyrG* marker. | This work | 14.3 | - |
| pYFAC | AMA1-based backbone vector with *pyrG* marker. | Chooi et al[3] | 14.2 | 168982 |
| pYFAC-ubi-Y-pyrG-mCherry | Expression of mCherry reporter from an AMA1-based vector with *ubi-Y-pyrG* marker. | This work | 16.4 | - |
| pYFAC-ubi-M-pyrG-mCherry | Expression of mCherry reporter from an AMA1-based vector with *ubi-M-pyrG* marker. | This work | 16.4 | - |
| pYFAC-ODC-pyrG-mCherry | Expression of mCherry reporter from an AMA1-based vector with *ODC-pyrG* marker. | This work | 16.2 | - |
| pYFAC-mCherry | Expression of mCherry reporter from an AMA1-based vector with *pyrG* marker. | This work | 16.1 | - |
| pYFAC-ubi-Y-pyrG-crRNAarraymicA | Expression of crRNA array targeting chromosomal *micA* for CRISPRa, from a AMA1- | This work | 15.8 | - |

| | | | | |
|---|---|---|---|---|
| | based vector with *ubi-Y-pyrG* marker. | | | |
| pYFAC-ubi-M-pyrG-crRNAarraymicA | Expression of crRNA array targeting chromosomal *micA* for CRISPRa, from a AMA1-based vector with *ubi-M-pyrG* marker. | This work | 15.8 | - |
| pYFAC-ODC-pyrG-crRNAarraymicA | Expression of crRNA array targeting chromosomal *micA* for CRISPRa, from a AMA1-based vector with ODC-*pyrG* marker. | This work | 15.5 | - |
| pYFAC-PYRG-crRNAarraymicA (pCRI040) | Expression of crRNA array targeting chromosomal *micA* for CRISPRa, from a AMA1-based vector with *pyrG* marker. | Roux et al 2020[4] | 15.5 | - |
| pYFAC-ubi-Y-pyrG-micA | Expression of the NRPS-like gene *micA* from a AMA1-based vector with *ubi-Y-pyrG* marker. | This work | 18.8 | - |
| pYFAC-ubi-M-pyrG-micA | Expression of the NRPS-like gene *micA* from a AMA1-based vector with *ubi-M-pyrG* marker. | This work | 18.8 | - |
| pYFAC-pyrG-micA | Expression of the NRPS-like gene *micA* from a AMA1-based vector with *pyrG* marker. | Roux et al 2020[4] | 18.6 | - |
| pYFAC-ubi-Y-CH2 | Expression of heterologous genes under alcohol induction with *ubi-Y-pyrG* marker. Promoters PalcA- | This work | 15.8 | 221131 |

| | | | | |
|---|---|---|---|---|
| | T1-PalcS/M-T2-PaldA expression cassette; unique restriction site: PacI for *PalcA*, NotI for *PalcS*, AsiSI for *PalcM*, AscI for *PaldA*. | | | |
| pYFAC-ubi-M-CH2 | Expression of heterologous genes under alcohol induction with *ubi-M-pyrG* marker. Promoters PalcA-T1-PalcS/M-T2-PaldA expression cassette; unique restriction site: PacI for *PalcA*, NotI for *PalcS*, AsiSI for *PalcM*, AscI for *PaldA*. | This work | 15.8 | 221132 |
| pYFAC-CH2 | Expression of heterologous genes under alcohol induction using the promoters PalcA-T1-PalcS/M-T2-PaldA expression cassette; unique restriction site: PacI for *PalcA*, NotI for *PalcS*, AsiSI for *PalcM*, AscI for *PaldA*. | [6] | 15.5 | 168978 |
| pYFAC-ubi-Y-CH2-neaC-neaD | Alcohol inducible expression of *neaC* and *neaD*. | This work | 20.5 | |
| pYFAC-ubi-M-CH2-neaC-neaD | Alcohol inducible expression of *neaC* and *neaD*. | This work | 20.5 | |
| pYFAC-CH2-neaC-neaD | Alcohol inducible expression of *neaC* and *neaD*. | This work | 20.2 | |

**Table S2 Oligonucleotides used in this study**

| Oligonuclotide | Sequence (5′ to 3′) | Purpose |
|---|---|---|
| pyrGRV | CGTGGGAATGGAGGGTTATATGC | Building pYFAC-ubi-Y-pyrG, pYFAC-ubi-M-pyrG, pYFAC-ODC-pyrG |
| pyrg_ubi_Fw | TATAACCCTCCATTCCCACGATGCAGATTTTCGTCAAGACTTTGAC | Building pYFAC-ubi-Y-pyrG and pYFAC-ubi-M-pyrG |
| PyrG_ubi_rv | GTCAATTGCGACTTGGACGACGTATTGGGCGCCAGGG | Building pYFAC-ubi-Y-pyrG and pYFAC-ubi-M-pyrG |
| PyrgCDS_F | TCGTCCAAGTCGCAATTGACTTA | |
| Pyrg-ODC-FW | ACAAGAAAGCATAACCTCCCTTTACAAAAAAGCCGGCTCCTCGTCCAAGTCGCAATTGAC | Building pYFAC-ODC-pyrG |
| pyrg-ODC-RV | GAGGTTATGCTTTCTTGTGCGCAGCTCATTGGAAGCATCGTGGGAATGGAGGGTTATATG | Building pYFAC-ODC-pyrG |
| PyrgPartR | GCCATATAAGCTTCCCAGCCT | |
| PKW-NotI-PacI-TrpcT | AACACAGTGGAGGACATACCCGTAATTTTCTGGGCGGCCGCTATTCGTTAATTAACGTCATGCATTGCAGATGAG | Amplifying expression cassette for crRNA array or mCherry for homology based cloning in pYFAC vector |
| PKW-GPda-Fw | CTTAGTAACCTCGCGGGTGTTCTTGACGATGGCATCCTGCCATGCGGAGAGACGGACGGT | Amplifying expression cassette for crRNA array or mCherry for homology based cloning in pYFAC vector |
| Pkw-micA-Fw | CGCGGGTGTTCTTGACGATGGCATCCTGCGGGGTACTGTTTCTGAATCCACTCTACATTC | Amplifying *micA* expression cassette |
| PyrG-micA-Rv | CAGTGGAGGACATACCCGTAATTTTCTGGGCGGCCGCTCCAATAGCTACGCACCAAACG | Amplifying *micA* expression cassette |
| FRR2249_scf5_8617(neaD)fg1CH2_NotI_F | TGAGATACCAAAGCATTGAGCCCAGAAACAGCAGAAGCatggcgatactatcgggc | Neoaspergillic acid cloning |
| FRR2249_scf5_8617(neaD)fg1CH2_NotI_R | AGTCTAAAGGTCTACAATCAATTCAGGCCGTATTCAGGGCctaaccctcttgacgcatc | Neoaspergillic acid cloning |
| FRR2249_scf5_8618(neaC)fg1CH2_PacI_F | TTAATTAGAACTCTTCCAATCCTATCACCTCGCCTTAatgtctggatttcccgatc | Neoaspergillic acid cloning |

| | | |
|---|---|---|
| FRR2249_scf5_8618(neaC)fg1CH2_PacI_R | CGCGCTCCACGGGGACTCGCTTCAATTTGTTCCGCTTAATtcaccctctagcaggcac | Neoaspergillic acid cloning |
| fragment_TpyrGoverhang_ampCENUra3_F | GATCACATCGGCAATTCTGATAACAGATGGTGCCATTGTCCCGTAATCAATTGCCCATTG | To build pYFAC-ubi-Y-CH2 and pYFAC-ubi-M-CH2 |
| fragment_TpyrGoverhang_ampCENUra3_R | GGCGAACGTGGCGAGAAAGG | To build pYFAC-ubi-Y-CH2 and pYFAC-ubi-M-CH2 |
| fragment_Alcs_F | ACGATGAGCCGCTCTTGCATC | To build pYFAC-ubi-Y-CH2 and pYFAC-ubi-M-CH2 |
| pKW-PacI-alcA-R | CACAGTGGAGGACATACCCGTAATTTTCTGGGCTTAATTAAGGCGAGGTGATAGGATTGG | To build pYFAC-ubi-Y-CH2 and pYFAC-ubi-M-CH2 |
| pyrG-PaldH-R | CTTCAACACAGTGGAGGACATACCCGTAATTTTCTGGGCTGATCTTCTCATCACGCCTCT | To build pYFAC-ubi-Y-CH2 and pYFAC-ubi-M-CH2 |
| PpyrG_PaldHoverhang_F | TGAAAATTACGAACCAAGAGGCGTGATGAGAAGATCAGCCCAGAAAATTACGGGTATGTC | To build pYFAC-ubi-Y-CH2 and pYFAC-ubi-M-CH2 |
| PpyrG_TalcAoverhang_F | TAGAACTCTTCCAATCCTATCACCTCGCCTTAATTAAGCCCAGAAAATTACGGGTATGTC | To build pYFAC-ubi-Y-CH2 and pYFAC-ubi-M-CH2 |
| mAmpR_R | CAGTGCTGCAATGATACCGC | To build pYFAC-ubi-Y-CH2 and pYFAC-ubi-M-CH2 |

**Table S3 *Aspergillus nidulans* strains built in this study**

| Strain number | *A. nidulans* parental strain genotype | Vector transformed |
|---|---|---|
| 1 | LO8030 [5] | pYFAC |
| 2 | LO8030 [5] | pYFAC-ubi-Y-pyrG-mCherry |
| 3 | LO8030 [5] | pYFAC-ubi-M-pyrG-mCherry |
| 4 | LO8030 [5] | pYFAC-ODC-pyrG-mCherry |
| 5 | LO8030 [5] | pYFAC-mCherry |
| 6 | LO8030 ; *PgpdA-dCas12aVPR-TtrpC*::ΔSterigmatocystin cluster(*AN7804–AN7825*) [4] | pYFAC-ubi-Y-pyrG-crRNAarraymicA |
| 7 | LO8030 ; *PgpdA-dCas12aVPR-TtrpC*::ΔSterigmatocystin cluster(*AN7804–AN7825*) [4] | pYFAC-ubi-M-pyrG-crRNAarraymicA |
| 8 | LO8030 ; *PgpdA-dCas12aVPR-TtrpC*::ΔSterigmatocystin cluster(*AN7804–AN7825*) [4] | pYFAC-ODC-pyrG-crRNAarraymicA |
| 9 | LO8030 ; *PgpdA-dCas12aVPR-TtrpC*::ΔSterigmatocystin cluster(*AN7804–AN7825*) [4] | pYFAC-PYRG-crRNAarraymicA (pCRI040) |
| 10 | LO8030 [5] | pYFAC-ubi-Y-pyrG-micA |
| 11 | LO8030 [5] | pYFAC-ubi-M-pyrG-micA |
| 12 | LO8030 [5] | pYFAC- pyrG-micA |
| 13 | LO8030 [5] | pYFAC-CH2 |
| 14 | LO8030 [5] | pYFAC-CH2-neaC-neaD |
| 15 | LO8030 [5] | pYFAC-ubi-Y-CH2-neaC-neaD |
| 16 | LO8030 [5] | pYFAC-ubi-M-CH2-neaC-neaD |

**Table S4 NMR data for neoaspergillic acid (600 ($^1$H) and 150 ($^{13}$C) MHz, $CDCl_3$).**

| pos | $\delta_C$, type | $\delta_H$ | gHMBC |
|---|---|---|---|
| 1 | 152.7, C | | |
| 2 | 152.2, C | | |
| 3 | 123.3, CH | 7.31 (s) | 1, 2, 4 |
| 4 | 136.4, C | | |
| 5 | 36.9, CH2 | 2.64, d (7.4) | 3, 4, 6, 7, 8 |
| 6 | 26.9, CH | 2.21, m | 4, 5, 7, 8 |
| 7 | 22.5, CH3 | 0.95, d (6.7) | 5, 6, 8 |
| 8 | 22.5, CH3 | 0.95, d (6.7) | 5, 6, 7 |
| 9 | 41.6, CH2 | 2.74, d (7.4) | 1, 2, 10, 11, 12 |
| 10 | 27.3, CH | 2.16, m | 2, 9, 11, 12 |

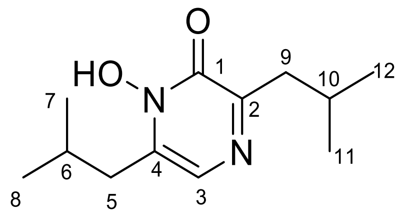

### Supporting text 1. Nucleotide sequence of degron tagged *pyrG* markers

| **Genetic part colour code** | **Description** | **Source** |
|---|---|---|
| AfpyrG exon | Auxotrophy marker | (Chooi et al., 2017) |
| AfpyrG intron | Auxotrophy marker | (Chooi et al., 2017) |
| Ubiquitin | Ubi-N degron | (Houser et al., 2012). |
| **Methionine codon** | Ubi-N degron | (Houser et al., 2012). |
| **Tyrosine codon** | Ubi-N degron | (Houser et al., 2012). |
| Δk sequence flexible linker | Ubi-N degron | (Houser et al., 2012). |
| ODCdegron | ODC degron | (Pohl, 2020). |

>UBI-M-pyrG_ORF
ATGCAGATTTTCGTCAAGACTTTGACCGGTAAAACCATAACATTGGAAGTTGAATCTTCC
GATACCATCGACAACGTTAAGTCGAAAATTCAAGACAAGGAAGGTATCCCTCCAGATCA
ACAAAGATTGATCTTTGCCGGTAAGCAGCTAGAAGACGGTAGAACGCTGTCTGATTACA
ACATTCAGAAGGAGTCCACCTTACATCTTGTGCTAAGGCTAAGAGGTGGT**ATG**CACGG
ATCCGGAGCTTGGCTGTTGCCCGTCTCACTGGTGAAAAGAAAAACCACCCTGGCGCCC
AATACGTCGTCCAAGTCGCAATTGACTTACGGTGCTCGAGCCAGCAAGCACCCCAATC
CTCTGGCAAAGAGACTTTTTGAGATTGCCGAAGCAAAGAAGACAAACGTTACCGTCTCT
GCTGATGTGACGACAACCCGAGAACTCCTGGACCTCGCTGACCGTACGGAAGCTGTTG
GATCCAATACATATGCCGTCTAGCAATGGACTAATCAACTTTTGATGATACAGGTCTCG
GTCCCTACATCGCCGTCATCAAGACACACATCGACATCCTCACCGATTTCAGCGTCGAC
ACTATCAATGGCCTGAATGTGCTGGCTCAAAAGCACAACTTTTTGATCTTCGAGGACCG
CAAATTCATCGACATCGGCAATACCGTCCAGAAGCAATACCACGGCGGTGCTCTGAGG
ATCTCCGAATGGGCCCACATTATCAACTGCAGCGTTCTCCCTGGCGAGGGCATCGTCG
AGGCTCTGGCCCAGACCGCATCTGCGCAAGACTTCCCCTATGGTCCTGAGAGAGGACT
GTTGGTCCTGGCAGAGATGACCTCCAAAGGATCGCTGGCTACGGGCGAGTATACCAAG
GCATCGGTTGACTACGCTCGCAAATACAAGAACTTCGTTATGGGTTTCGTGTCGACGCG
GGCCCTGACGGAAGTGCAGTCGGATGTGTCTTCAGCCTCGGAGGATGAAGATTTCGTG
GTCTTCACGACGGGTGTGAACCTCTCTTCCAAAGGAGATAAGCTTGGACAGCAATACCA
GACTCCTGCATCGGCTATTGGACGCGGTGCCGACTTTATCATCGCCGGTCGAGGCATC
TACGCTGCTCCCGACCCGGTTGAAGCTGCACAGCGGTACCAGAAAGAAGGCTGGGAA
GCTTATATGGCCAGAGTATGCGGCAAGTCATGA

>UBI-Y-pyrG_ORF
ATGCAGATTTTCGTCAAGACTTTGACCGGTAAAACCATAACATTGGAAGTTGAATCTTCC
GATACCATCGACAACGTTAAGTCGAAAATTCAAGACAAGGAAGGTATCCCTCCAGATCA
ACAAAGATTGATCTTTGCCGGTAAGCAGCTAGAAGACGGTAGAACGCTGTCTGATTACA
ACATTCAGAAGGAGTCCACCTTACATCTTGTGCTAAGGCTAAGAGGTGGT**TAT**CACGGA
TCCGGAGCTTGGCTGTTGCCCGTCTCACTGGTGAAAAGAAAAACCACCCTGGCGCCCA
ATACGTCGTCCAAGTCGCAATTGACTTACGGTGCTCGAGCCAGCAAGCACCCCAATCC
TCTGGCAAAGAGACTTTTTGAGATTGCCGAAGCAAAGAAGACAAACGTTACCGTCTCTG
CTGATGTGACGACAACCCGAGAACTCCTGGACCTCGCTGACCGTACGGAAGCTGTTGG
ATCCAATACATATGCCGTCTAGCAATGGACTAATCAACTTTTGATGATACAGGTCTCGGT
CCCTACATCGCCGTCATCAAGACACACATCGACATCCTCACCGATTTCAGCGTCGACAC
TATCAATGGCCTGAATGTGCTGGCTCAAAAGCACAACTTTTTGATCTTCGAGGACCGCA
AATTCATCGACATCGGCAATACCGTCCAGAAGCAATACCACGGCGGTGCTCTGAGGAT
CTCCGAATGGGCCCACATTATCAACTGCAGCGTTCTCCCTGGCGAGGGCATCGTCGAG
GCTCTGGCCCAGACCGCATCTGCGCAAGACTTCCCCTATGGTCCTGAGAGAGGACTGT
TGGTCCTGGCAGAGATGACCTCCAAAGGATCGCTGGCTACGGGCGAGTATACCAAGG
CATCGGTTGACTACGCTCGCAAATACAAGAACTTCGTTATGGGTTTCGTGTCGACGCGG

GCCCTGACGGAAGTGCAGTCGGATGTGTCTTCAGCCTCGGAGGATGAAGATTTCGTGG
TCTTCACGACGGGTGTGAACCTCTCTTCCAAAGGAGATAAGCTTGGACAGCAATACCAG
ACTCCTGCATCGGCTATTGGACGCGGTGCCGACTTTATCATCGCCGGTCGAGGCATCT
ACGCTGCTCCCGACCCGGTTGAAGCTGCACAGCGGTACCAGAAAGAAGGCTGGGAAG
CTTATATGGCCAGAGTATGCGGCAAGTCATGA

>ODC-pyrG_ORF
ATGCTTCCAATGAGCTGCGCACAAGAAAGCATAACCTCCCTTTACAAAAAAGCCGGCTC
CTCGTCCAAGTCGCAATTGACTTACGGTGCTCGAGCCAGCAAGCACCCCAATCCTCTG
GCAAAGAGACTTTTTGAGATTGCCGAAGCAAAGAAGACAAACGTTACCGTCTCTGCTGA
TGTGACGACAACCCGAGAACTCCTGGACCTCGCTGACCGTACGGAAGCTGTTGGATCC
AATACATATGCCGTCTAGCAATGGACTAATCAACTTTTGATGATACAGGTCTCGGTCCCT
ACATCGCCGTCATCAAGACACACATCGACATCCTCACCGATTTCAGCGTCGACACTATC
AATGGCCTGAATGTGCTGGCTCAAAAGCACAACTTTTTGATCTTCGAGGACCGCAAATT
CATCGACATCGGCAATACCGTCCAGAAGCAATACCACGGCGGTGCTCTGAGGATCTCC
GAATGGGCCCACATTATCAACTGCAGCGTTCTCCCTGGCGAGGGCATCGTCGAGGCTC
TGGCCCAGACCGCATCTGCGCAAGACTTCCCCTATGGTCCTGAGAGAGGACTGTTGGT
CCTGGCAGAGATGACCTCCAAAGGATCGCTGGCTACGGGCGAGTATACCAAGGCATC
GGTTGACTACGCTCGCAAATACAAGAACTTCGTTATGGGTTTCGTGTCGACGCGGGCC
CTGACGGAAGTGCAGTCGGATGTGTCTTCAGCCTCGGAGGATGAAGATTTCGTGGTCT
TCACGACGGGTGTGAACCTCTCTTCCAAAGGAGATAAGCTTGGACAGCAATACCAGAC
TCCTGCATCGGCTATTGGACGCGGTGCCGACTTTATCATCGCCGGTCGAGGCATCTAC
GCTGCTCCCGACCCGGTTGAAGCTGCACAGCGGTACCAGAAAGAAGGCTGGGAAGCT
TATATGGCCAGAGTATGCGGCAAGTCATGA
